## Supplementary material for "Identification of a Binding Site for Small Molecule Inhibitors Targeting Human TRPM4": Cryo-EM data collection, refinement, and validation statistics for SMA solubilized HsTRPM4 samples.

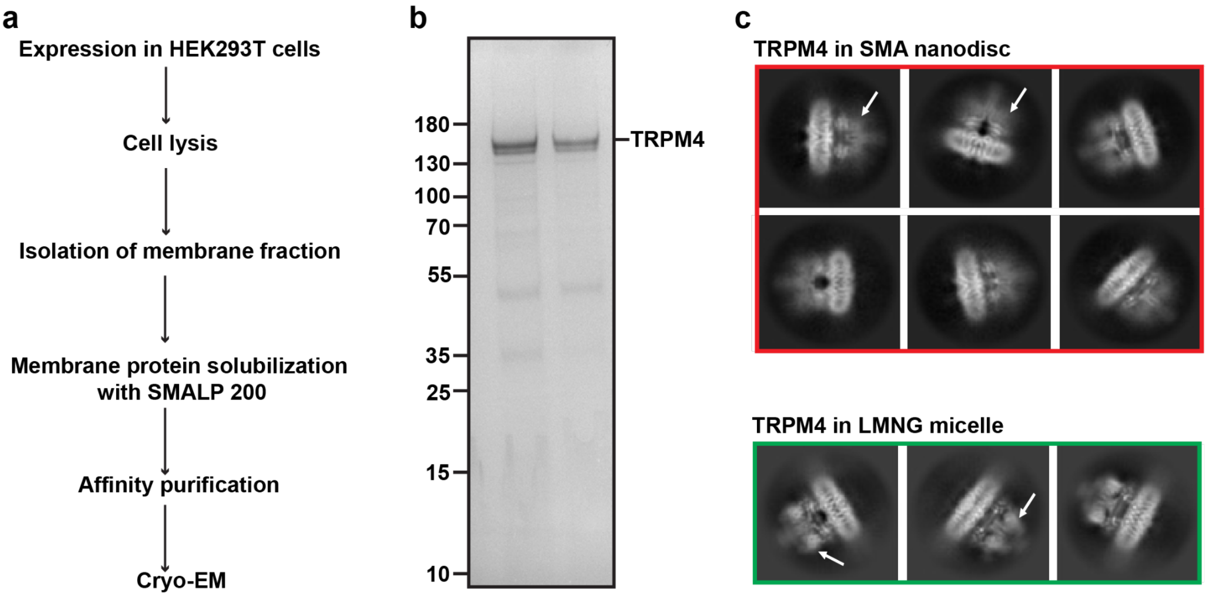


**Supplementary Fig. 1**: Purification of TRPM4 in SMA-extracted native lipid nanodiscs. (a) Biochemical workflow for isolating and purifying HsTRPM4 in native lipid nanodiscs using SMA (SMALP-200). (b) SDS-PAGE gel of purified HsTRPM4 following affinity purification. (c) Cryo-EM 2D class averages of TRPM4 in SMA-extracted native lipid nanodiscs and LMNG detergent micelles. White arrows indicate the positions of MHR1/2 in the TRPM4 2D classes.

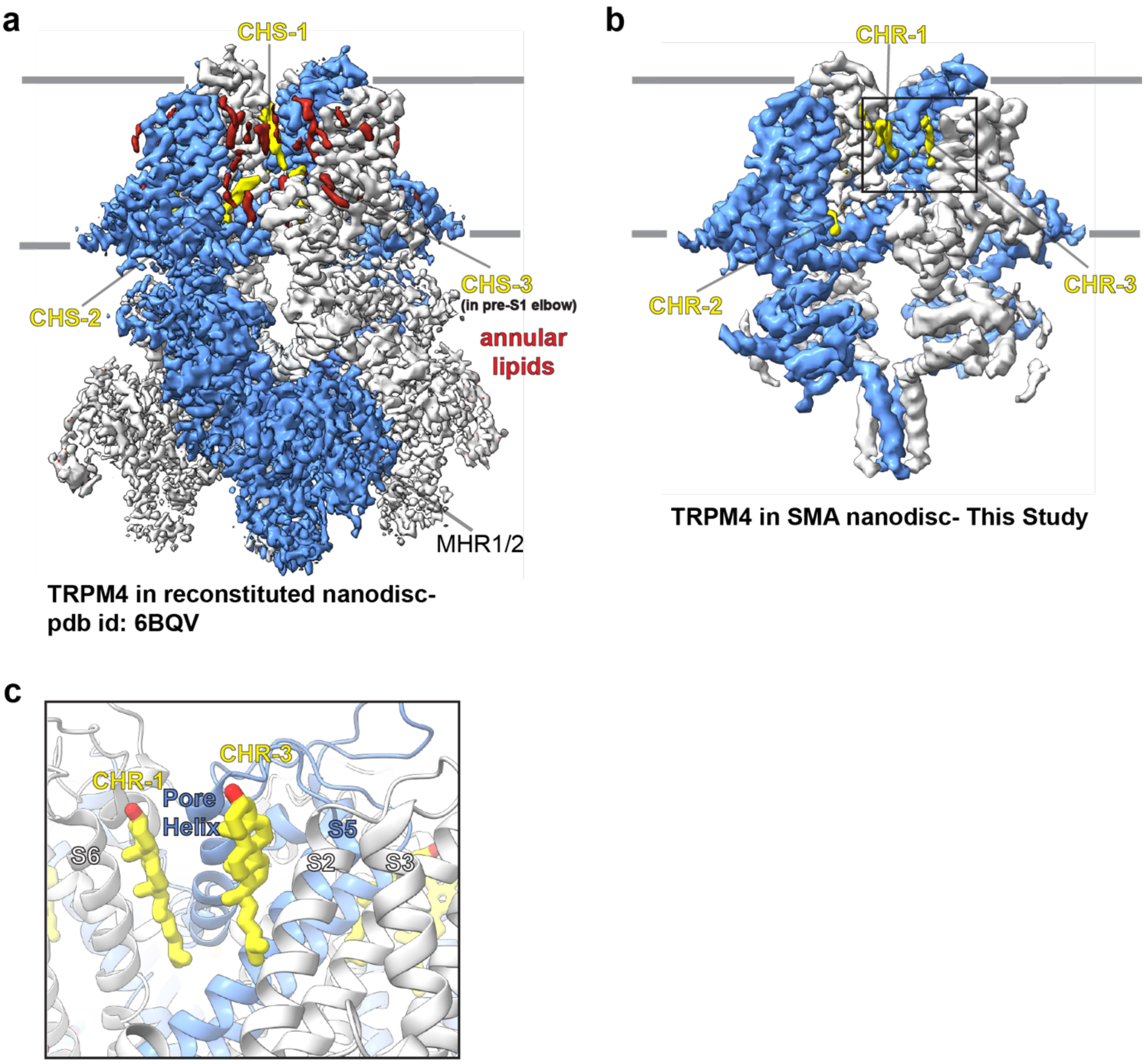


**Supplementary Fig. 2**: Comparison of the annular lipid environment between the cryo-EM maps of HsTRPM4 in reconstituted lipid nanodiscs (PDB 6BQV) [26] (a) and SMA-extracted native lipid nanodiscs (b). CHS represents cholesteryl hemisuccinate in (a), and CHR represents cholesterol in (b). (c) The location of CHR-1 and CHR-3 is shown from the cryo-EM structure of HsTRPM4 in native lipid nanodiscs.

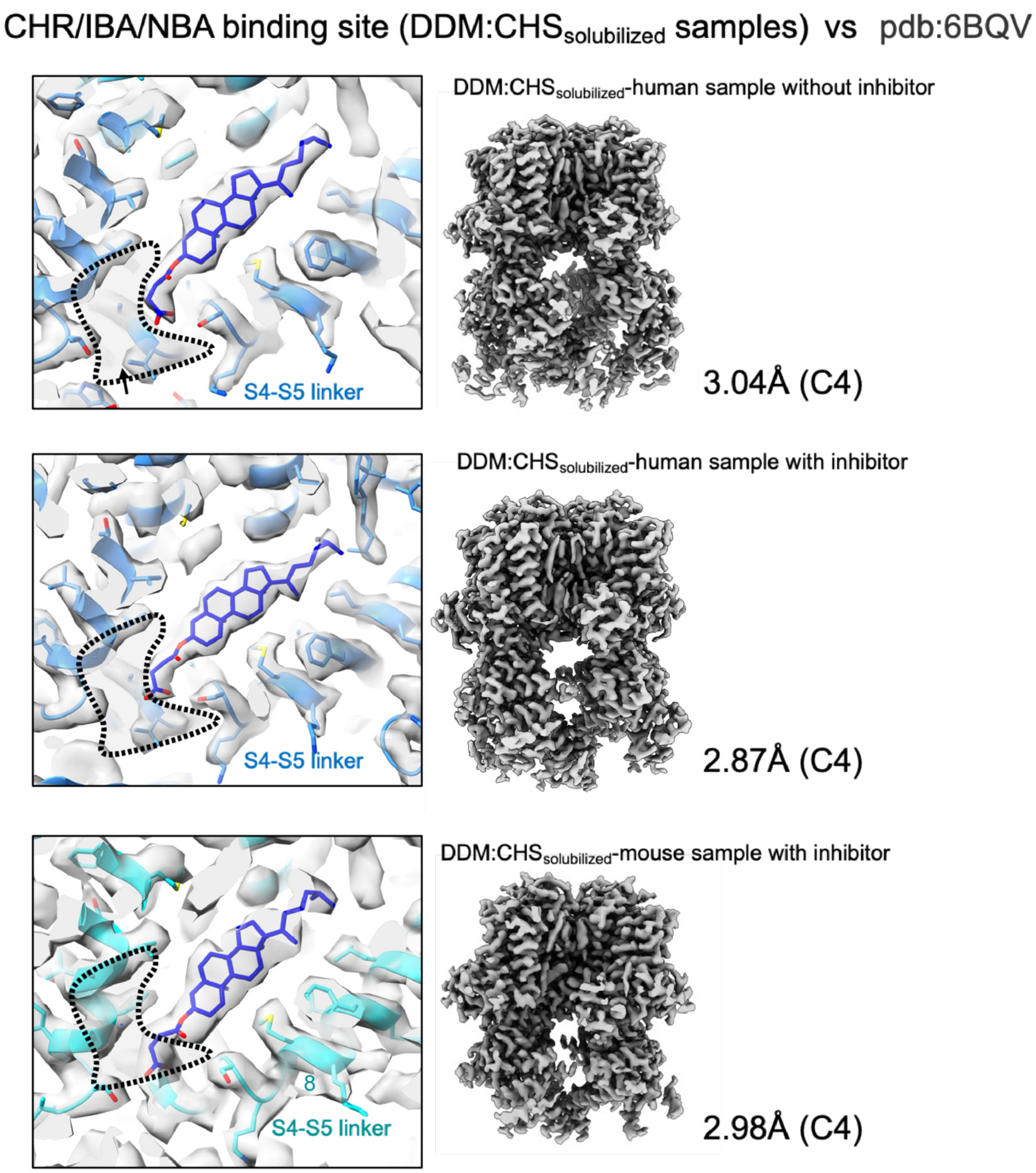


**Supplementary Fig. 3**: Cryo-EM maps of detergent-solubilized TRPM4 generated in this study highlight the presence of CHS in the drug binding site when fitted with the published model (PBD 6BQV). The dashed line indicates the location of the drug-binding site determined in this study.

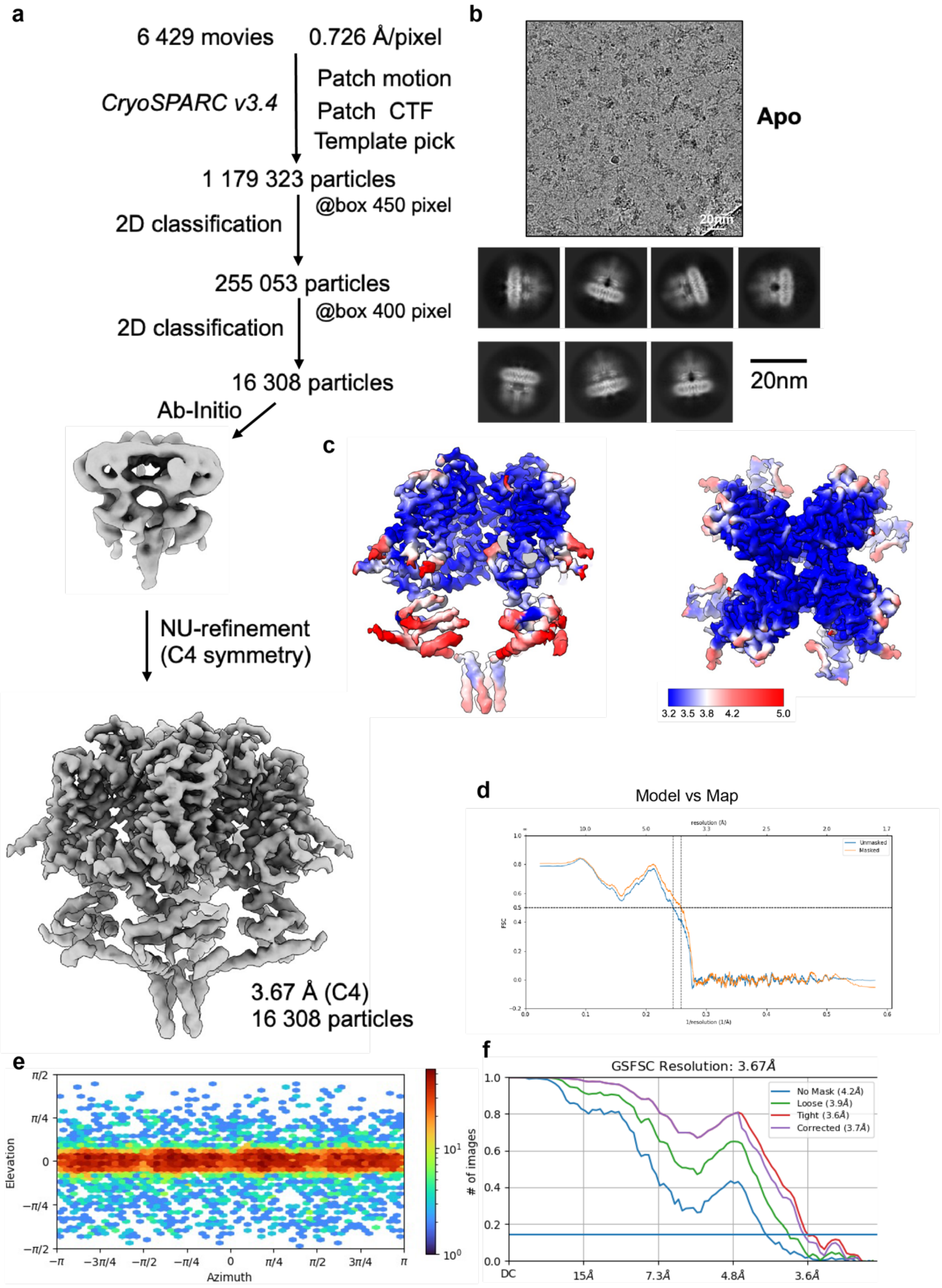


**Supplementary Fig. 4**: Cryo-EM data processing workflow and map resolution (a) The image processing workflow of $HsTRPM4_{apo}$. (b) micrograph and 2D classes (c) Local resolution (d) Model vs map FSC curves. (e) particle direction distribution. (f) FSC curves.

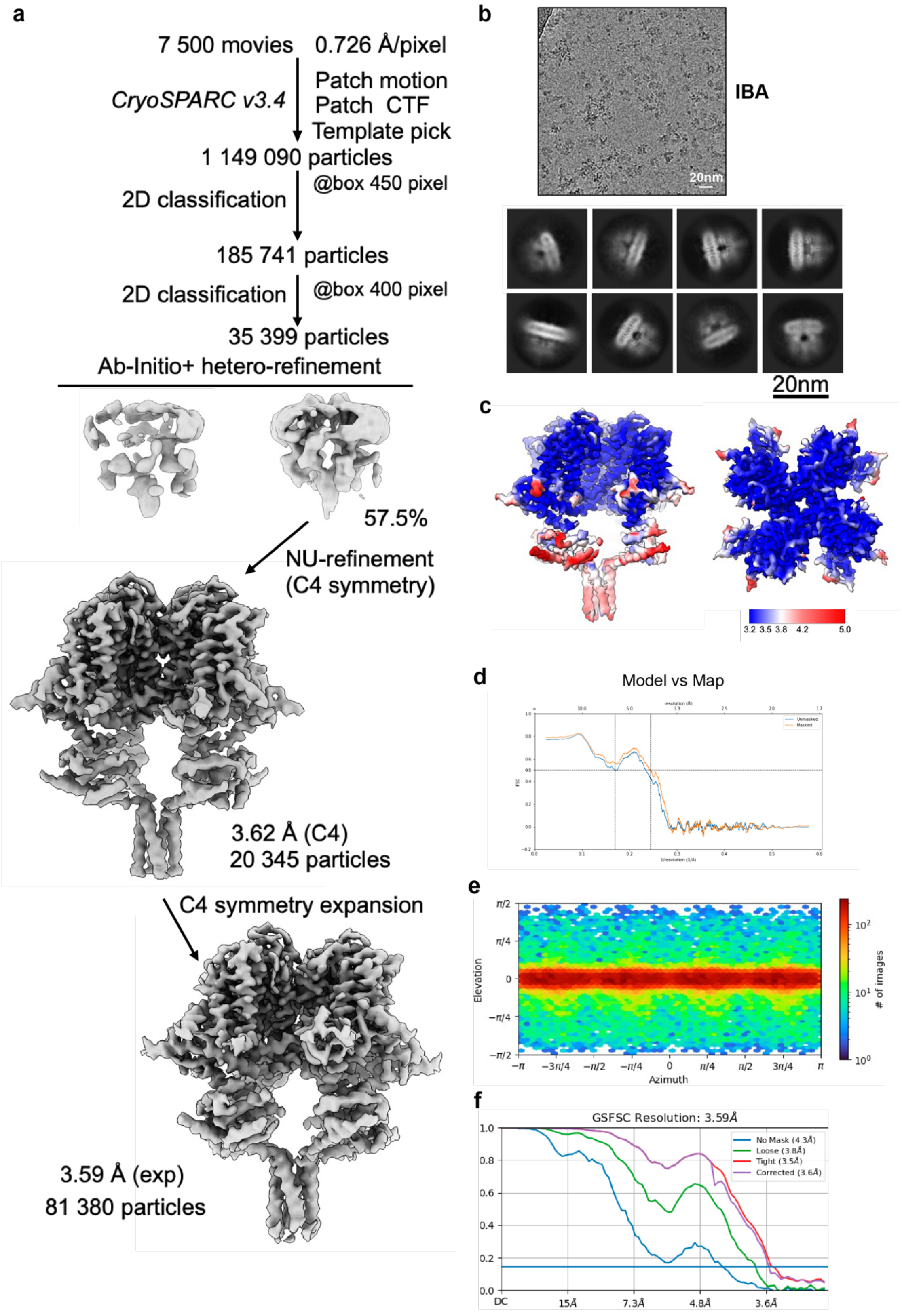


**Supplementary Fig. 5**: Cryo-EM data processing workflow and 3D reconstructions (a) The image processing workflow of $HsTRPM4_{IBA}$. (b) Micrograph and 2D classes. (c) Local resolution. (d) Model vs map FSC curves. (e) Particle direction distribution. (f) FSC curves.

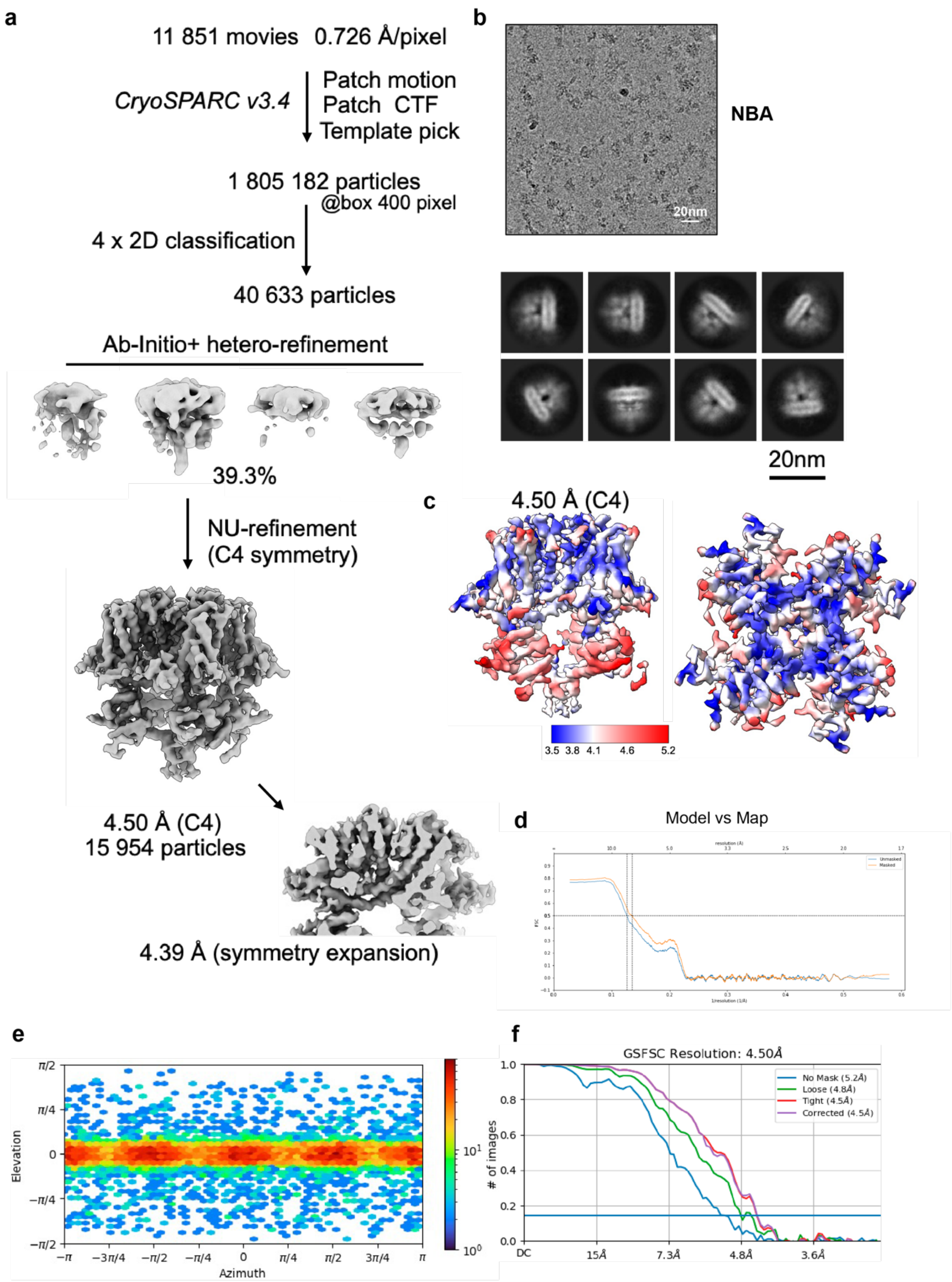


**Supplementary Fig. 6**: Cryo-EM data processing workflow and 3D reconstructions. (a) The image processing workflow of $HsTRPM4_{NBA}$. (b) Micrograph and 2D classes. (c) Local resolution. (d) Model vs. map FSC curves. (e) Particle direction distribution. (f) FSC curves.

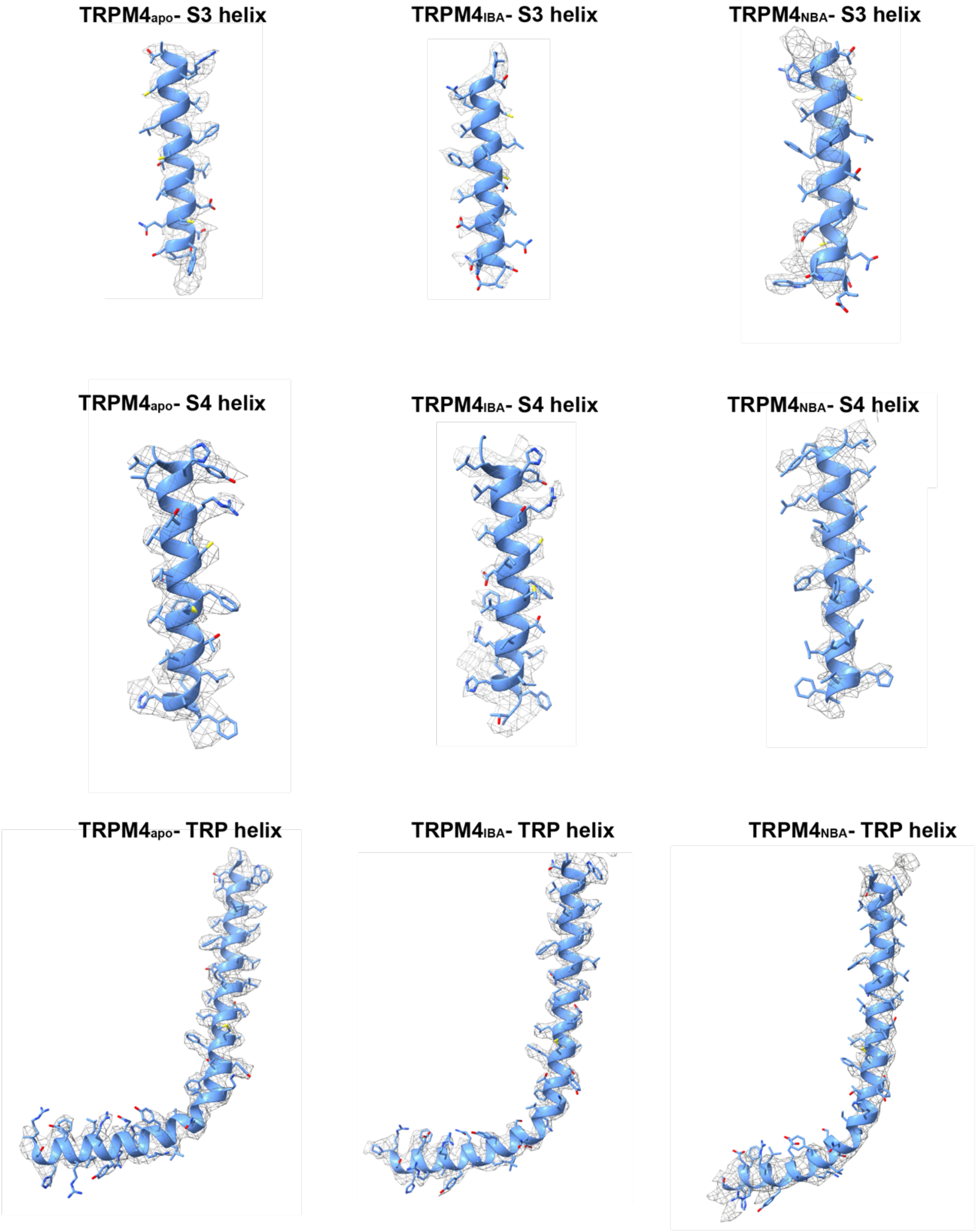


**Supplementary Fig. 7**: Zoom-in view of the cryo-EM maps of structural elements HsTRPM4$_{apo}$, HsTRPM4$_{IBA}$ and HsTRPM4$_{NBA}$.

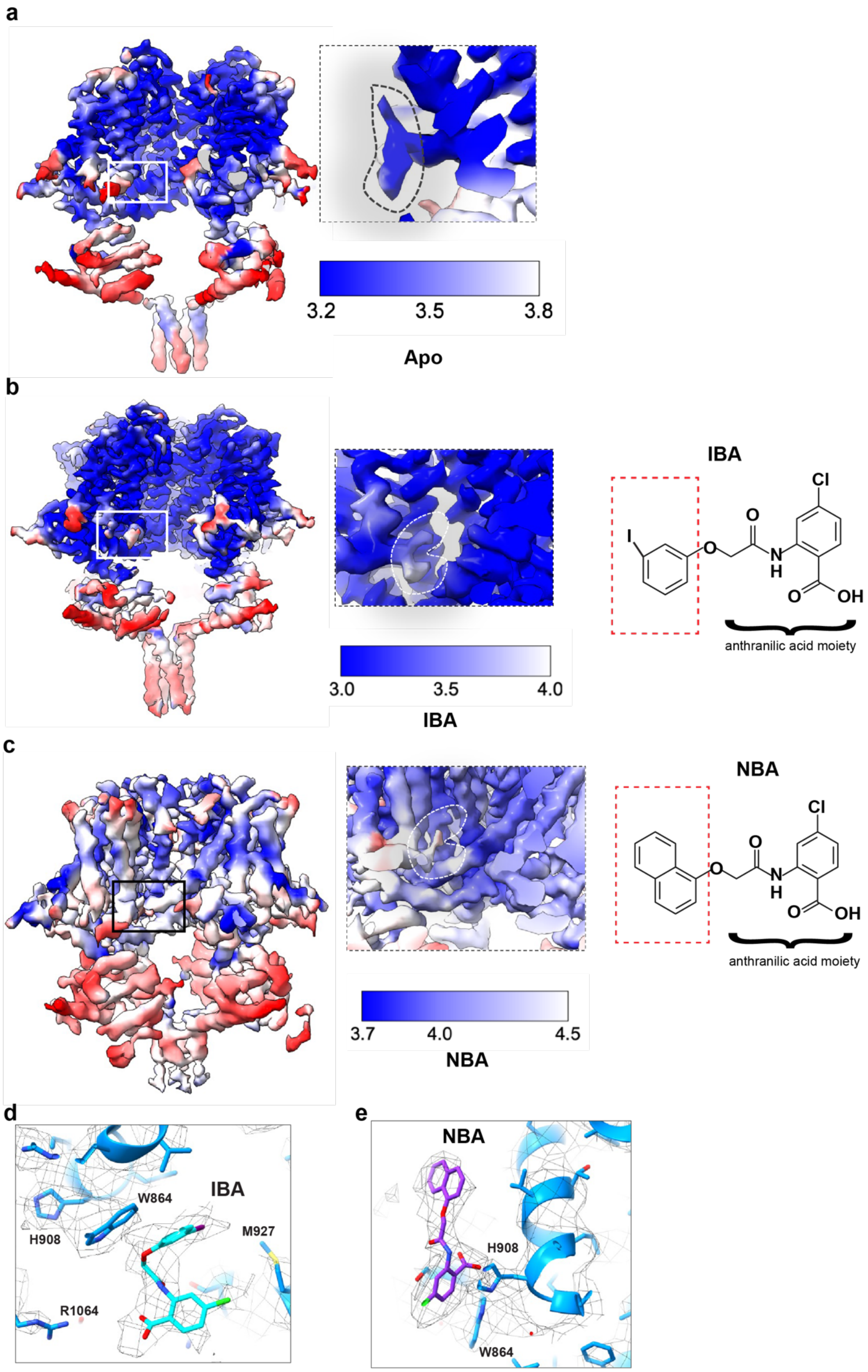


**Supplementary Fig. 8**: Local resolution maps of SMA solubilized HsTRPM4 generated in this study. (a) Map for the $HsTRPM4_{apo}$ with the density contained in the drug binding site highlighted. (b) Map for $HsTRPM4_{IBA}$ with the density contained in the drug binding site highlighted. The chemical structure of IBA is shown on the right. (c) Map for $HsTRPM4_{NBA}$ with the density contained in the drug binding site highlighted. The chemical structure of the NBA is shown on the right. (d and e) Zoom-in of NBA and IBA with the cryo-EM density in mesh representation in different views compared to Figure 2 e and f.

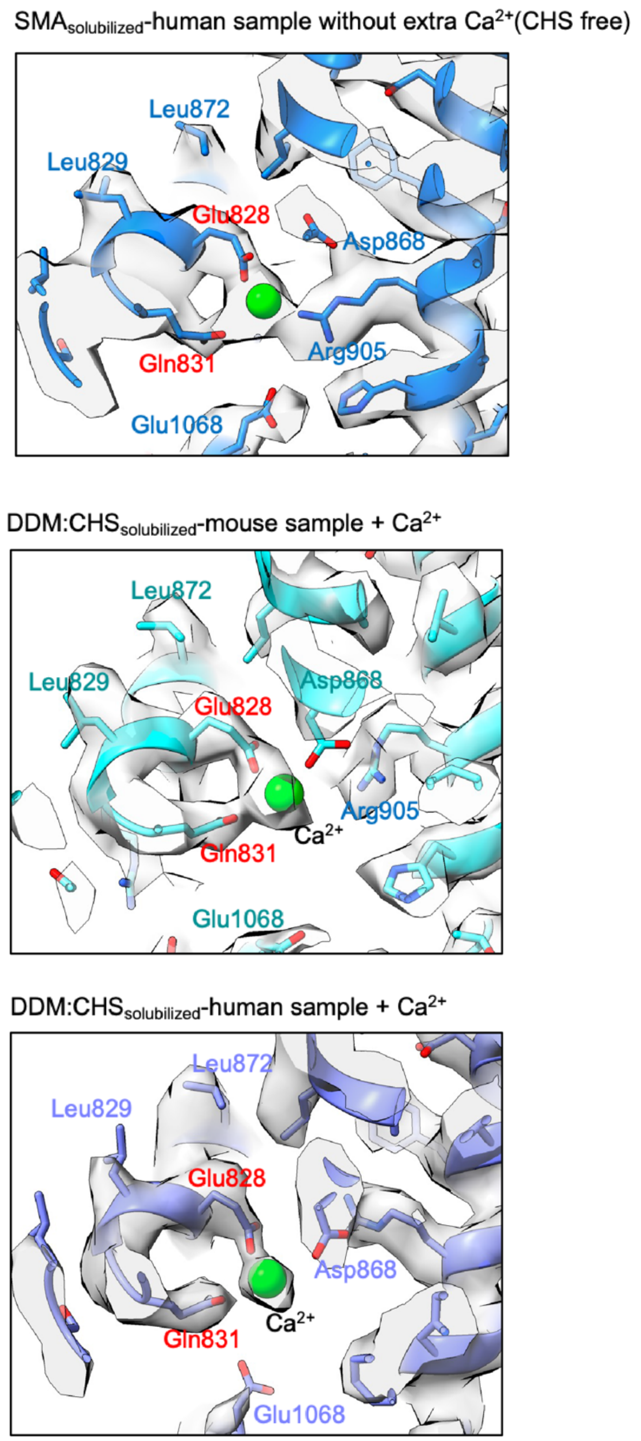


**Supplementary Fig. 9**: Identification of cryo-EM densities for $Ca^{2+}$ ions in the maps generated in this study. The published HsTRPM4 model (pdb-id: 6BQV) is fitted into each map. Clear density is observed which could be putatively occupied by a $Ca^{2+}$ in the map for the SMA solubilized human TRPM4 sample without the addition of extra calcium, which is comparable to the density shown for detergent-solubilized mouse and human TRPM4 with calcium added to the sample.

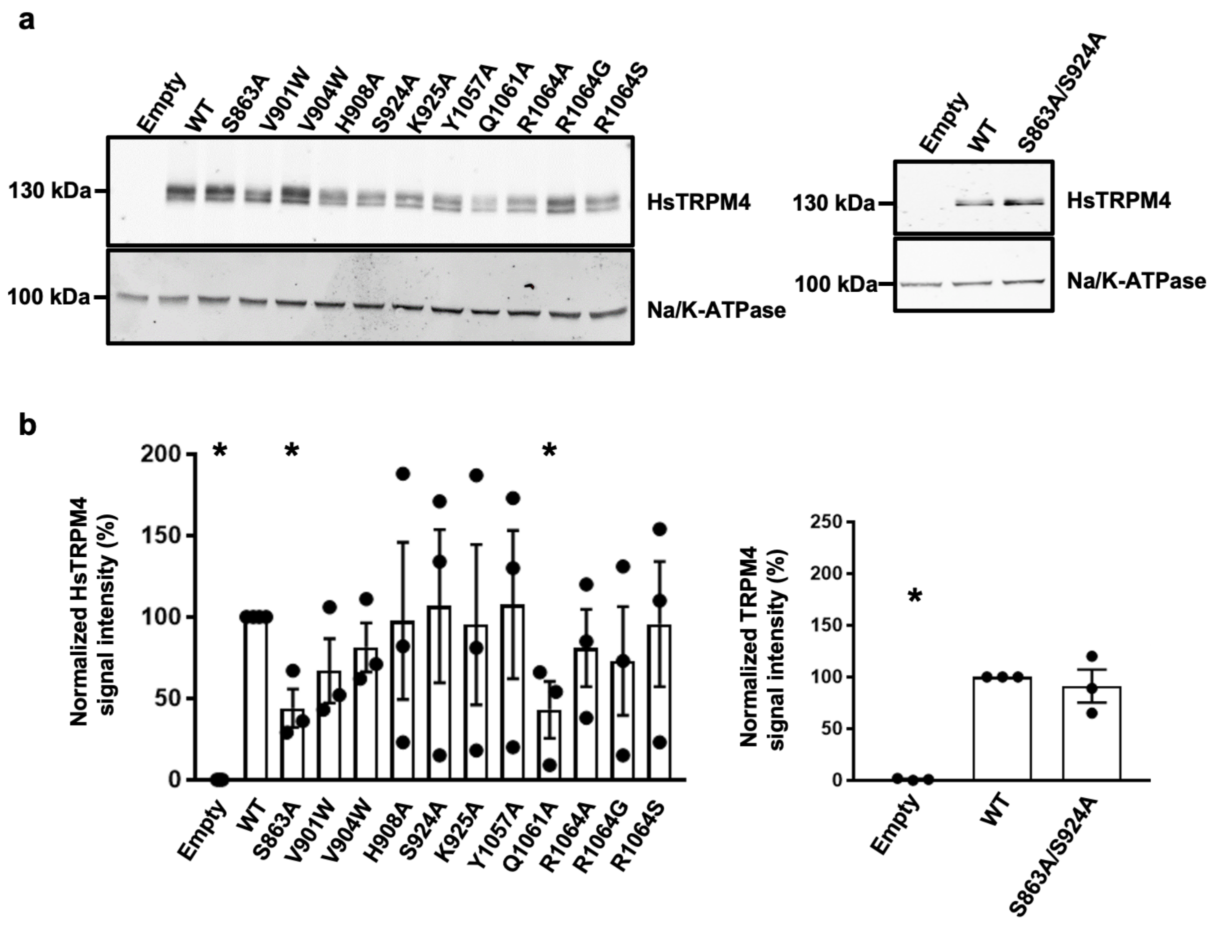


**Supplementary Fig. 10**: Expression of HsTRPM4 variants. (a) Western blot showing the expression of wildtype (WT) and HsTRPM4 variants expressed in HEK293 cells. (b) Graphical plot of relative western blot intensities between wildtype (WT) and HsTRPM4 variants. (*) represents the p-value <0.05 compared to the wildtype condition with n=3.

a

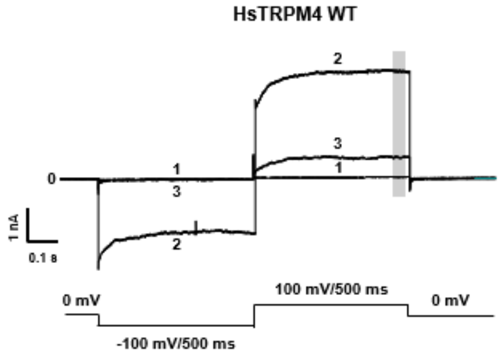


b

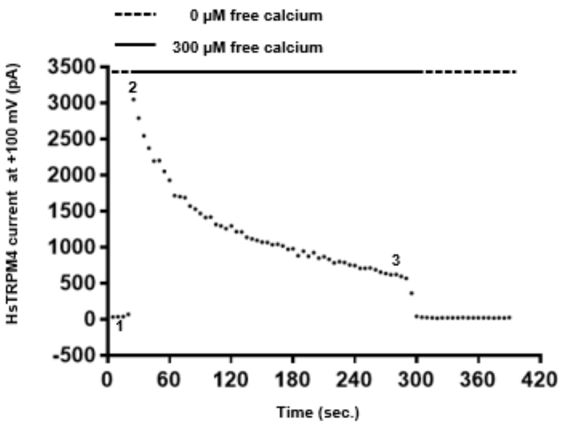


**Supplementary Fig. 11**: Time course of wildtype calcium-activated sodium current for HsTRPM4. (a) Example of raw traces (currents) from three different time points 1, 2, and 3 from the time course shown in (b) recorded from one cell expressing HsTRPM4 channels. Without free calcium (time point 1 in (b)), no calcium-activated sodium current is recordable at -100 mV or +100 mV (trace n°1 in (a)). Perfusion of 300 µM of free calcium concentration leads to the rapid activation of the channel (time point 2 in (b)), and a calcium-activated sodium current is recordable (trace n°2 in (a)) at both voltages (-100 mV and +100 mV). Following this quick activation, a rapid run-down of the calcium-activated sodium current mediated by the HsTRPM4 channel, due to the depletion of the PIP2 element, is observable until the current reaches stability (time point 3 in (b)). After the run-down, the calcium-activated sodium current amplitude is quantified by calculating the amplitude of this current during the last 100 ms at +100 mV (grey zone in (a)). As depicted in trace number 3 in (a), the run-down is generally more pronounced at -100 mV, leading to almost a complete inhibition of the calcium-activated sodium current at this voltage.

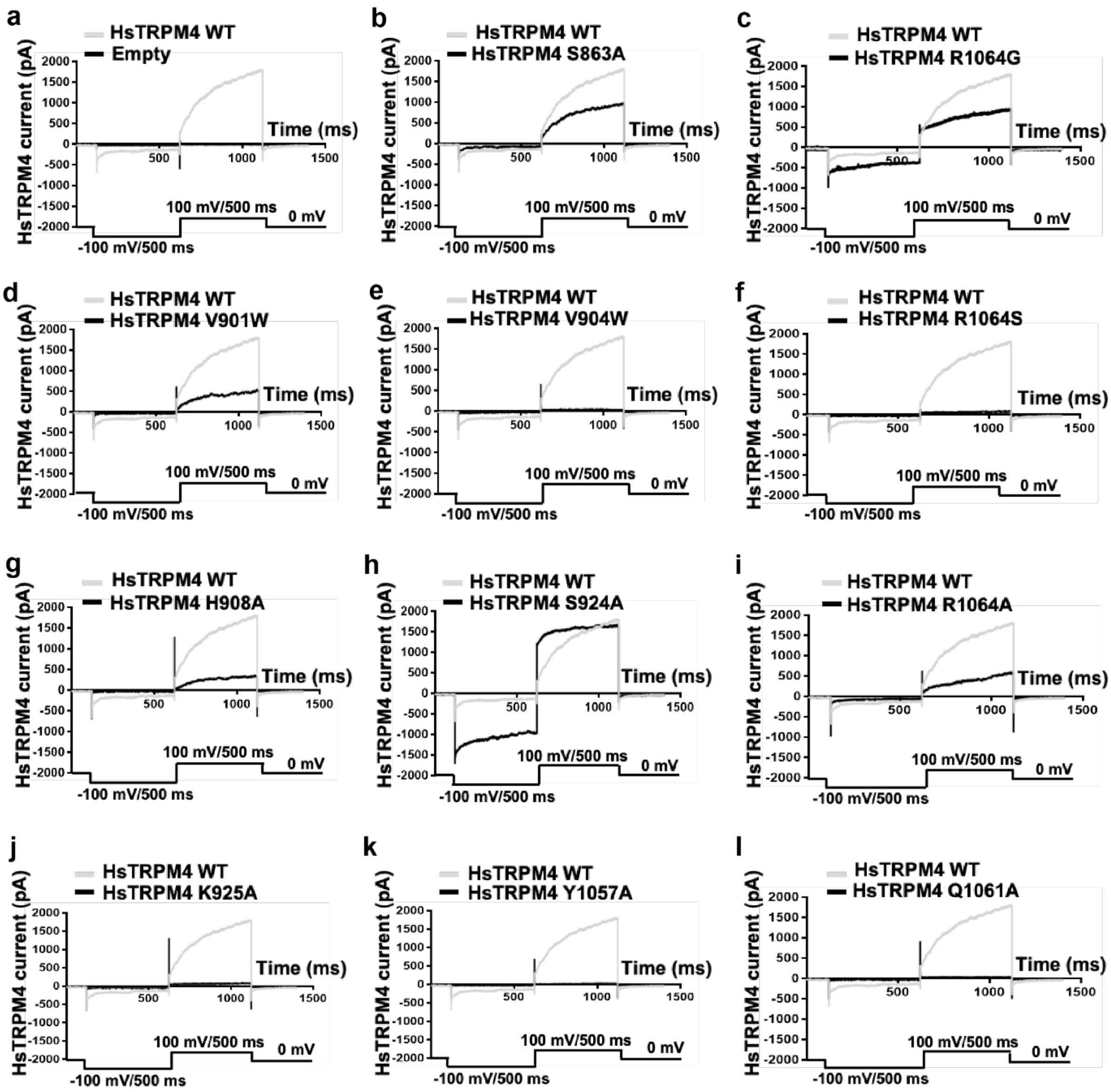


**Supplementary Fig. 12**: Calcium-activated sodium currents for HsTRPM4. (a to l) Representative traces of wildtype (WT, gray traces) and variants HsTRPM4 currents (black traces): S863A, V901W, V904W, H908A, S924A, K925A, Y1057A, Q1061A, R1064A, R1064G, and R1064S. Calcium-activated sodium currents for HsTRPM4 variant channels are generally smaller than WT currents, as depicted in Figure 3e, due to the amino acid alteration.

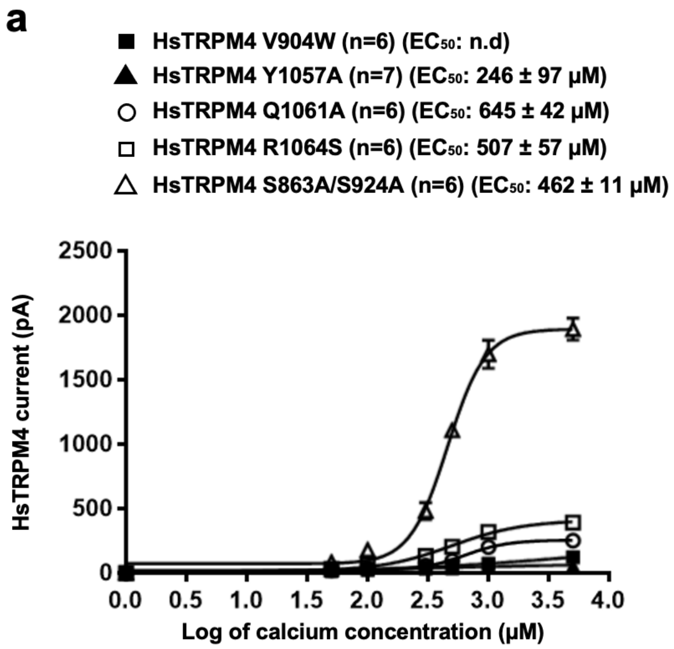


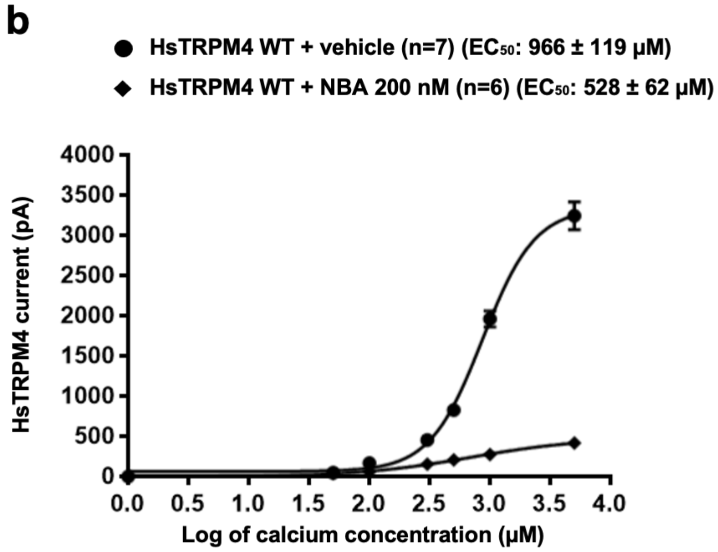


**Supplementary Fig. 13**: Calcium sensitivity curves of wildtype and variant HsTRPM4. (a) Calcium sensitivity curves of a few loss-of-function HsTRPM4 variants. (n): number of cells. n.d: not determinable. (b) Effect of NBA (200 nM) on the calcium sensitivity on wildtype (WT) HsTRPM4 channels.

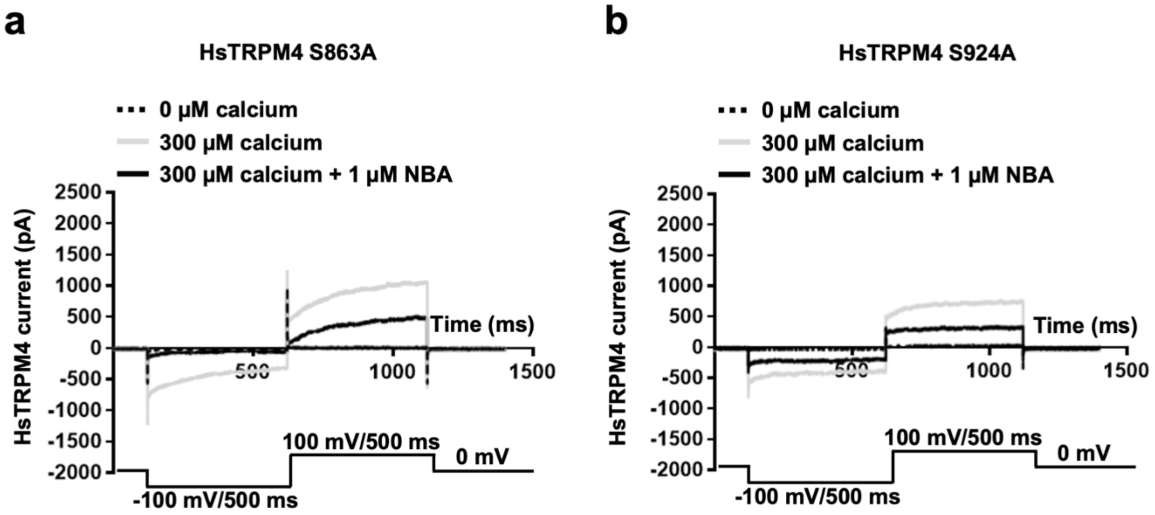


**Supplementary Fig. 14**: Calcium-activated sodium currents from HsTRPM4 channels. (a and b) Representative traces of variants HsTRPM4 current: S863A and S924A in the absence of free calcium in the solution (dotted black line), in the presence of 300 μM of free calcium concentration (grey line), and in the presence of 300 μM of free calcium concentration and 1 μM NBA (black line).
